# Virus-host interactome and proteomic survey of PMBCs from COVID-19 patients reveal potential virulence factors influencing SARS-CoV-2 pathogenesis

**DOI:** 10.1101/2020.03.31.019216

**Authors:** Jingjiao Li, Mingquan Guo, Xiaoxu Tian, Chengrong Liu, Xin Wang, Xing Yang, Ping Wu, Zixuan Xiao, Yafei Qu, Yue Yin, Joyce Fu, Zhaoqin Zhu, Zhenshan Liu, Chao Peng, Tongyu Zhu, Qiming Liang

## Abstract

The ongoing coronavirus disease (COVID-19) pandemic caused by Severe Acute Respiratory Syndrome Coronavirus-2 (SARS-CoV-2) is a global public health concern due to relatively easy person-to-person transmission and the current lack of effective antiviral therapy. However, the exact molecular mechanisms of SARS-CoV-2 pathogenesis remain largely unknown. We exploited an integrated proteomics approach to systematically investigate intra-viral and virus-host interactomes for the identification of unrealized SARS-CoV-2 host targets and participation of cellular proteins in the response to viral infection using peripheral blood mononuclear cells (PBMCs) isolated from COVID-19 patients. Using this approach, we elucidated 251 host proteins targeted by SARS-CoV-2 and more than 200 host proteins that are significantly perturbed in COVID-19 derived PBMCs. From the interactome, we further identified that non-structural protein nsp9 and nsp10 interact with NKRF, a NF-КB repressor, and may precipitate the strong IL-8/IL-6 mediated chemotaxis of neutrophils and overexuberant host inflammatory response observed in COVID-19 patients. Our integrative study not only presents a systematic examination of SARS-CoV-2-induced perturbation of host targets and cellular networks to reflect disease etiology, but also reveals insights into the mechanisms by which SARS-CoV-2 triggers cytokine storms and represents a powerful resource in the quest for therapeutic intervention.

## MAIN TEXT

In December 2019, Severe Acute Respiratory Syndrome Coronavirus-2 [SARS-CoV-2, previously known by the provisional name ‘2019 novel coronavirus’ (2019-nCoV) given by the World Health Organization (WHO)], was first isolated from a cluster of patients with pneumonia. This novel strain of coronavirus has been identified as the causative agent for the epidemic of acute respiratory distress syndrome [also referred to as coronavirus disease 2019 (COVID-19)] in Wuhan, China^1–3^. Since then, the COVID-19 outbreak has rapidly swept all the provinces of China and has disseminated globally to 202 countries and territories through traveling^4^. On 31 January 2020, the WHO declared SARS-CoV-2 a Public Health Emergency of International Concern^5^. As of 27 March 2020, the current pandemic of SARS-CoV-2 has caused 512,701 confirmed infection cases including 23,495 deaths in 202 countries and territories, calling for extended efforts to understand the molecular drivers of its immune evasion and pathogenesis.

COVID-19 is a potential zoonotic disease with low to moderate mortality rate (2-5%) and is primarily transmitted through droplet and direct contact with infected persons or incubation carriers^6^. SARS-CoV-2 has a preferential tropism for airway epithelial cells mediated through the accessibility of ACE2 receptor^7–10^. The common clinical manifestations of fever and cough in conjunction with laboratory results of progressively increasing neutrophil counts and leukocytopenia in serum samples taken from COVID-19 patients at various stages of the illness indicates uncontrolled neutrophil extravasation and activation^11,12^. Neutrophils are the most abundant type of white blood cells in human blood and serve as the first responders in the line of defense against acute infection. Chemical signals such as Interleukin-8 (IL-8) and IL-6 stimulate neutrophil chemotaxis and recruitment to the site of infection where they congregate and cause the release of more proinflammatory mediators that in turn activate additional inflammatory signals, leading to cytokine storms that may progress to vital organ failure^13^. Indeed, a large number of macrophages and neutrophils were found in alveolar cavities in COVID-19 fatal patients^14^, suggesting their roles for cytokine storms. While uncontrolled neutrophil activity and aberrant cytokine response contributes to SARS-CoV-2 pathogenesis in COVID-19, how SARS-CoV-2 causes these immunological changes, especially the enhancement of neutrophil chemotaxis and activation remains largely undetermined.

SARS-CoV-2 is a single-stranded positive-sense RNA virus and belongs to the β-coronavirus family consisting of many animal species and human pathogens including SARS-CoV and Middle East Respiratory Syndrome (MERS)-CoV. The ~30 kb non-segmented genome of SARS-CoV-2 encodes 14 major open reading frames (ORFs), which are further processed into 4 structural proteins [glycoprotein spike (S), membrane (M), envelope (E), and nucleocapsid (N)], 16 non-structural proteins (nsp1 to nsp16), and at least 8 accessory proteins (ORF3a, ORF6, ORF7a, ORF7b, ORF8, ORF9a, ORF9b, and ORF10)^15^. Similar to SARS-CoV, spike protein of SARS-CoV-2 directly interacts with human ACE2 expressed on human airway epithelia to facilitate virus entry^7–10^. However, host targets for other viral proteins remain largely unknown, which limits our understanding of SARS-CoV-2 immune evasion and pathogenesis, further impeding anti-viral drug development and design of therapeutic approaches.

### Genome-wide screens for SARS-CoV-2 intra-viral protein-protein interactions

Limited by their relatively small genomes, viral proteins form different combinations of protein complexes to fulfill a variety of critical functions during viral life cycle, making the investigation of intra-viral protein-protein interactions (PPIs) a requirement. To systematically characterize the intra-viral PPI networks, both genome-wide yeast-two hybrid (Y2H) screens and co-immunoprecipitations (co-IP) experiments with SARS-CoV-2 genes were performed. All the predicted viral genes were synthesized based on the SARS-CoV-2 Wuhan-Hu-1 isolate (GenBank: MN908947). The entire library was cloned into Y2H DNA-binding domain (pACT2) and activation domain (pGBTK7) vectors and then transformed into the Y2HGold yeast strain (Clontech). Nsp3 was truncated into two smaller fragments in this experiment since large genes tend to yield fewer interactions in the Y2H assay. All the possible pair-wise combinations were examined independently for the activation of the reporter genes HIS3 and ADE2 in triplicates. We captured 19 PPIs from the Y2H screens, including nsp15-nsp2, ORF7b-nsp1, N-nsp4, ORF3a-nsp4, ORF7b-nsp7, nsp7-nsp8, nsp12-nsp8, nsp7-nsp9, nsp14-nsp10, nsp16-nsp10, ORF6-nsp14, E-E, N-E, ORF3a-E, ORF9b-E, N-N, nsp7-ORF7a, nsp8-ORF8, and ORF9b-ORF9b (**Extended Data Fig.1**). The Co-IP assay resulted in the identification of 52 PPIs (**Extended Data Fig.1-2**). Some of the interactions between viral proteins, such as the association between nsp12 and nsp8, were detected by both Y2H and co-IP. Some viral proteins, such as M, N, E, nsp2, nsp5, nsp8, ORF6, ORF7a, ORF7b, ORF9b, and ORF10 displayed self-association, suggesting their dimer or oligomer forms may carry out important functions during infection. In total, we characterized 58 distinct intra-viral PPIs amongst 28 SARS-CoV-2 genes that may promote virus replication and can be expected to be important to immune evasion and viral pathogenesis. (**Fig.1a**).

Among the 4 classical structural proteins, 3 PPIs were detected (**Fig.1b**); these interactions of structural proteins may be critical in the organization of viral particles during viral replication and assembly. We also found 8 PPIs between structural proteins and accessory proteins (ORF3a, ORF6, ORF7a, ORF7b, ORF9b, and ORF10) and 6 PPIs between structural proteins and non-structural proteins (nsp2, nsp4, nsp5, nsp8, and nsp16) (**Fig. 1b**), These PPI may play a pivotal role in viral structural protein processing, modification, and trafficking, supporting their importance for viral particle assembly and egress.

**Figure 1.**
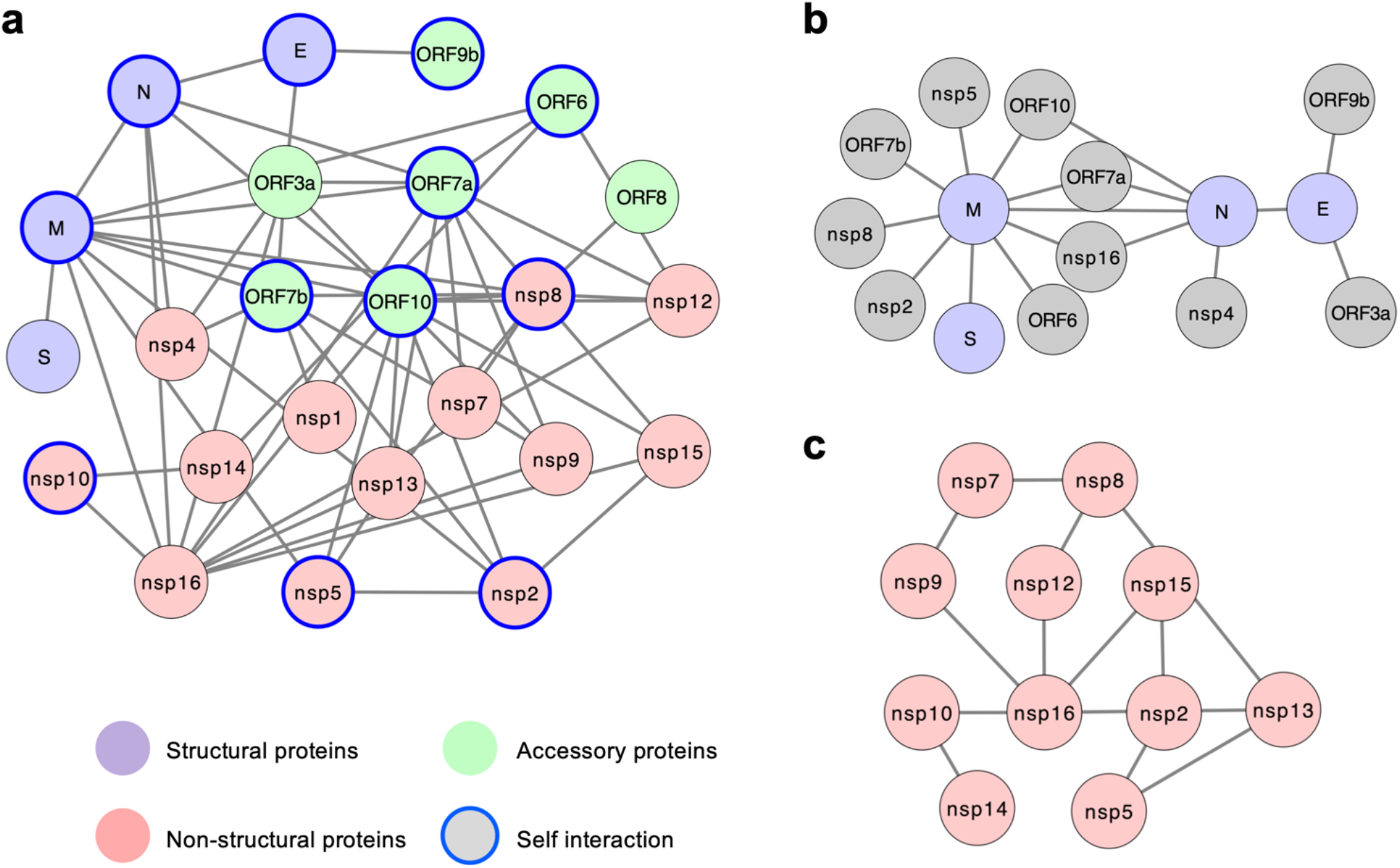
SARS-CoV-2 intra-viral protein-protein interaction network. (**a**) Viral interactions based on Y2H screens and co-IP experiments as described in Extended Data Fig. 1. Different types of viral proteins are labelled with the indicated colors. Self-associated viral proteins are circled in blue. (**b**) Interactions between viral structural proteins and other viral proteins. (**c**) Interaction network amongst viral non-structural proteins.

While structural proteins are components for the capsid and the envelope of the virus, several non-structural proteins (nsps), such as nsp3, nsp5, nsp12, nsp14, and nsp16 have associated enzymatic activities essential for viral RNA replication and viral protein translation. In addition, nsps can function as virulence factors that inhibit the host’s immune system defenses. We speculate that the relatively large number of nsp interactions detected (12) might imply a crucial role of nsps in the formation of multimeric complexes during the viral life cycle (**Fig.1c**). The SARS-CoV-2 ORF1ab (the precursor of all 16 nsps) shares 94.4% amino acid identity to SARS-CoV ORF1ab. A number of structural studies posits the methyltransferase activity of SARS-CoV nsp14 or nsp16 are facilitated by nsp10-nsp14 and nsp10-nsp16 associations, respectively^16,17^, which are also observed in SARS-CoV-2. Thus, our intra-viral PPI network provides the foundation to dissect the role of these viral proteins in mediating the synthesis, assembly and processing of viral proteins for both SARS-CoV-2 and the closely related SARS-CoV. Disruption of these intra-viral PPIs by repurposing small molecule or peptide drugs that target specific epitopes can be a treatment strategy against COVID-19^18^.

### Genome-wide proteomic screen for SARS-CoV-2 viral-host protein-protein interactions

To better understand the roles of these viral proteins during the virus life cycle, it is pivotal to establish the SARS-CoV-2-host interactome in human cells. We overexpressed the plasmids encoding each SARS-CoV-2 encoded gene with N-terminal 3xFlag-epitope in HEK293 cells and purified each SARS-CoV-2 protein complex by affinity purification. Co-purified cellular proteins were subsequently analyzed by liquid chromatography and tandem mass spectrometry (AP-LC-MS). From AP-LC-MS analysis, 251 cellular proteins were identified to interact with SARS-CoV-2 proteins, resulting in 631 high-confidence interactions (**Fig.2** and **Table 1**). We identified previously reported proteins that had bona fide associations with SARS-CoV proteins, such as ROA0 and subunits of the ATPase (AT1A1 and AT1A2)^19^, further substantiating these cellular processes as vital targets of the coronavirus family. SARS-CoV-2 viral proteins also interacts with the subunits of 40S ribosomal proteins (RS5, RS10, RS12, RS14, RS17, RS18, and RS27), T complex proteins (TCPA, TCPD, TCPE, TCPG, TCPH, TCPQ, and TCPZ), and proteasome-related proteins (PRS4, PRS7, PRS8, PSA, PSA4, PSA6, PSA7, PSD11, PSD13, PSMD1, PSMD2, PSMD6, and PSMD7). These viral-cellular protein interactions might have extensive cytopathic effects on the host cell, often altering host gene translation, protein folding, and degradation pathways during viral infection to assist viral growth and proliferation (**Extended Data Fig.3a-b)**. The virus may also co-opt the complex membranous network surrounding and residing within the host cell, such as the endoplasmic reticulum, Golgi apparatus, and plasma membranes, so long as the membrane systems possess host proteins that lead to a productive infection (**Fig.2**). Most importantly, the AP-LC-MS analysis uncovered host targets pertinent in inflammatory and innate immune responses, which may help explain the common respiratory symptoms. Key viral-cellular PPIs that may influence the pathogenesis of SARS-CoV-2 include interaction between NKRF (an endogenous repressor of IL-8/IL-6 synthesis) and nsp9/nsp10/nsp12/nsp13/nsp15^20,21^, C1QBP (involved in complement C1 activation) and nsp9/nsp10^22^, G3BP2 (anti-viral stress granule protein) and N^23^, and RAE1 (an interferon-inducible mRNA nuclear export protein) and ORF6^24^. These observations provide an overview of host proteins or pathways that are modulated by SARS-CoV-2 viral proteins and these PPIs might serve as potential targets for drug development or repurposing against COVID-19.

**Figure 2.**
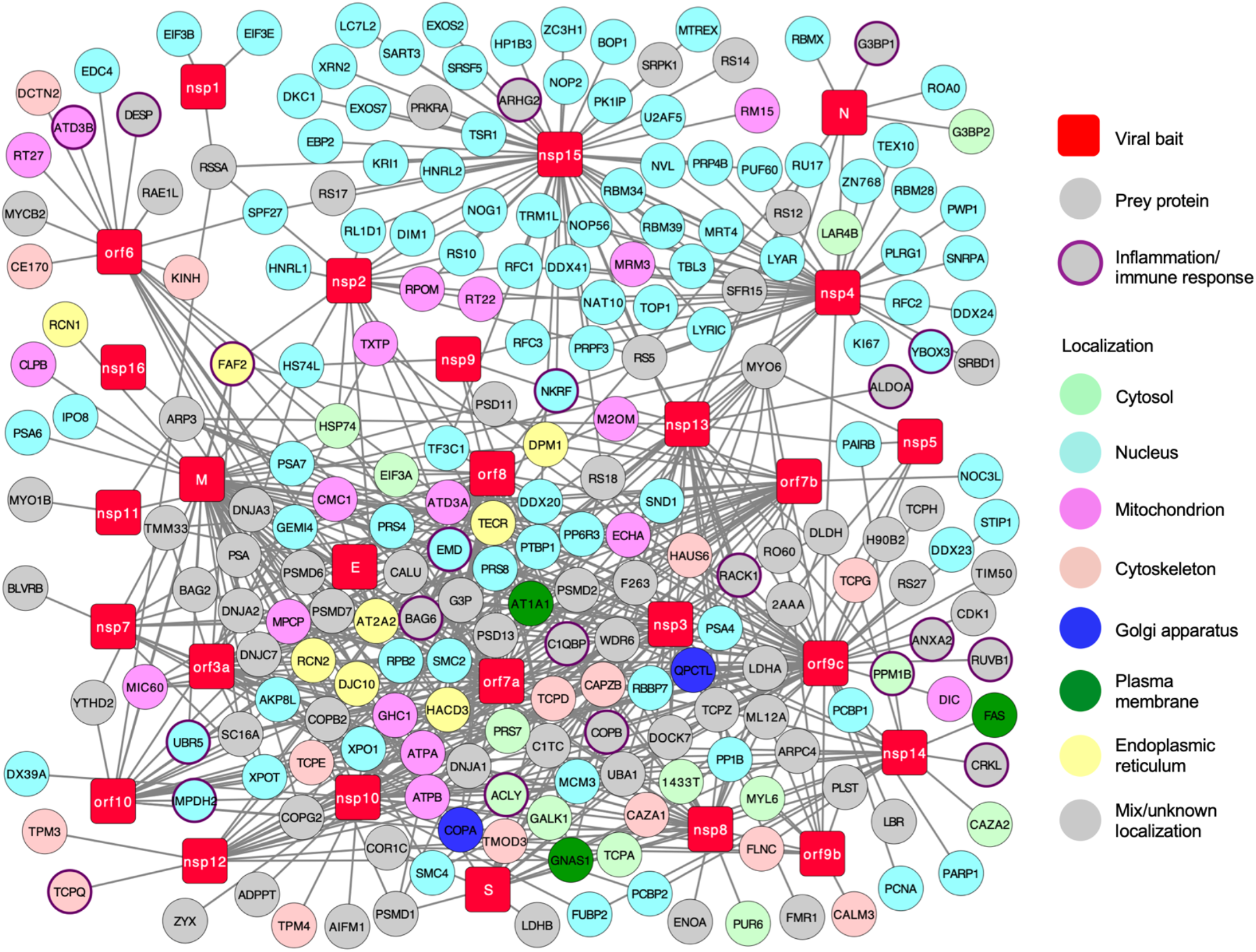
SARS-CoV-2-human protein-protein interaction network. Network representation of the high-confidence SARS-CoV-2-host interactome in HEK293 cells. There are 27 SARS-CoV-2 bait proteins (red squares) and 251 interacting host protein (circles). Subcellular localizations of host proteins are labelled with indicated colors. Proteins with known functions in inflammation or immune responses are circled in purple.

### Global proteome profiling of PBMCs from COVID-19 patients

To elucidate the mechanism and effect of SARS-CoV-2 infection in patients, we determined the global proteome profile changes in PBMCs samples taken from healthy donors compared with PBMCs collected from COVID-19 patients with mild or severe symptoms. The PBMC samples were obtained from the Shanghai Public Health Clinical Center and includes 6 healthy donors, 22 COVID-19 patients with mild symptoms (COVID-19-MS), and 13 COVID-19 patients with severe symptoms (COVID-19-SS) (**Extended Data Fig.4**). PBMCs were harvested from the leftover blood samples and all the patients were confirmed negative for influenza A virus. Following the Guideline of Novel Coronavirus Laboratory Safety (Edition 2), PBMCs were collected, lysed in denaturing buffer (see Methods) at the Shanghai Public Health Clinical Center and sent for proteome analysis at the National Facility for Protein Science in Shanghai.

A total of 6,739 proteins were identified in our study, of which 4,274 proteins could be quantified in all 26 PMBCs samples. Proteins with less than three quantitation values did not pass filtering threshold and were excluded from the quantitation and statistical analysis. 5,522 proteins were quantified according to criteria and included in global proteomic profiling to reveal a list of 220 proteins found to be differentially expressed [|log2 (fold change)| ≥ 1, n≥3] in the PBMCs of COVID-19-MS compared with that of healthy donors: 115 proteins showed a statistically significant increase while 105 proteins showed reduction in expression when comparing COVID-19-MS with control. (**Fig.3a**, **Extended Data Fig.5a-b and Table 2**). Functional analysis by DAVID and Cluster Profiler revealed that these genes were enriched in specific cellular biological processes, including positive regulation of neutrophil chemotaxis, neutrophil activation, type I interferon signaling pathway, inflammatory response, and antigen processing and presentation (**Fig.3b** and **Extended Data Fig.5c-d**), suggesting an active innate immune response against SARS-CoV-2 infection. Analysis of the global proteomic changes reveal significant upregulation of CXCR2, PRG3, LBP, MMP25, CRP, NLRP1 in the COVID-19-MS PBMC cohort (**Fig.3c**). CXCR2 is a subunit of IL-8 receptor and facilitates IL-8-induced neutrophil chemotaxis while PRG3 has been reported to stimulate neutrophil superoxide production and IL-8 release^25,26^. LBP positively regulates neutrophil response upon LPS stimulation and MMP25 inactivates alpha-1 proteinase inhibitor to facilitate the trans-endothelial migration of neutrophil to inflammatory sites during infection^27,28^. These differentially expressed proteins support a model by which SARS-CoV-2 triggers cytokine release, such as IL-8, in the early stage of infection, resulting in uncontrolled infiltration and activation of neutrophils to the sites of infection followed by eventual progression to cytokine storm and respiratory distress. Indeed, laboratory results reveal increased neutrophils to ~90% of the white blood cell count as COVID-19 progresses to critical stages^11,12^. The elevation of IL-8 and IL-6 but not IL-17 in PMBCs collected from COVID-19 patients indicates possible roles for IL-8/IL-6 in the severe inflammatory response induced by SARS-CoV-2 (**Fig.3d-e** and **Extended Data Fig.5e**).

**Figure 3.**
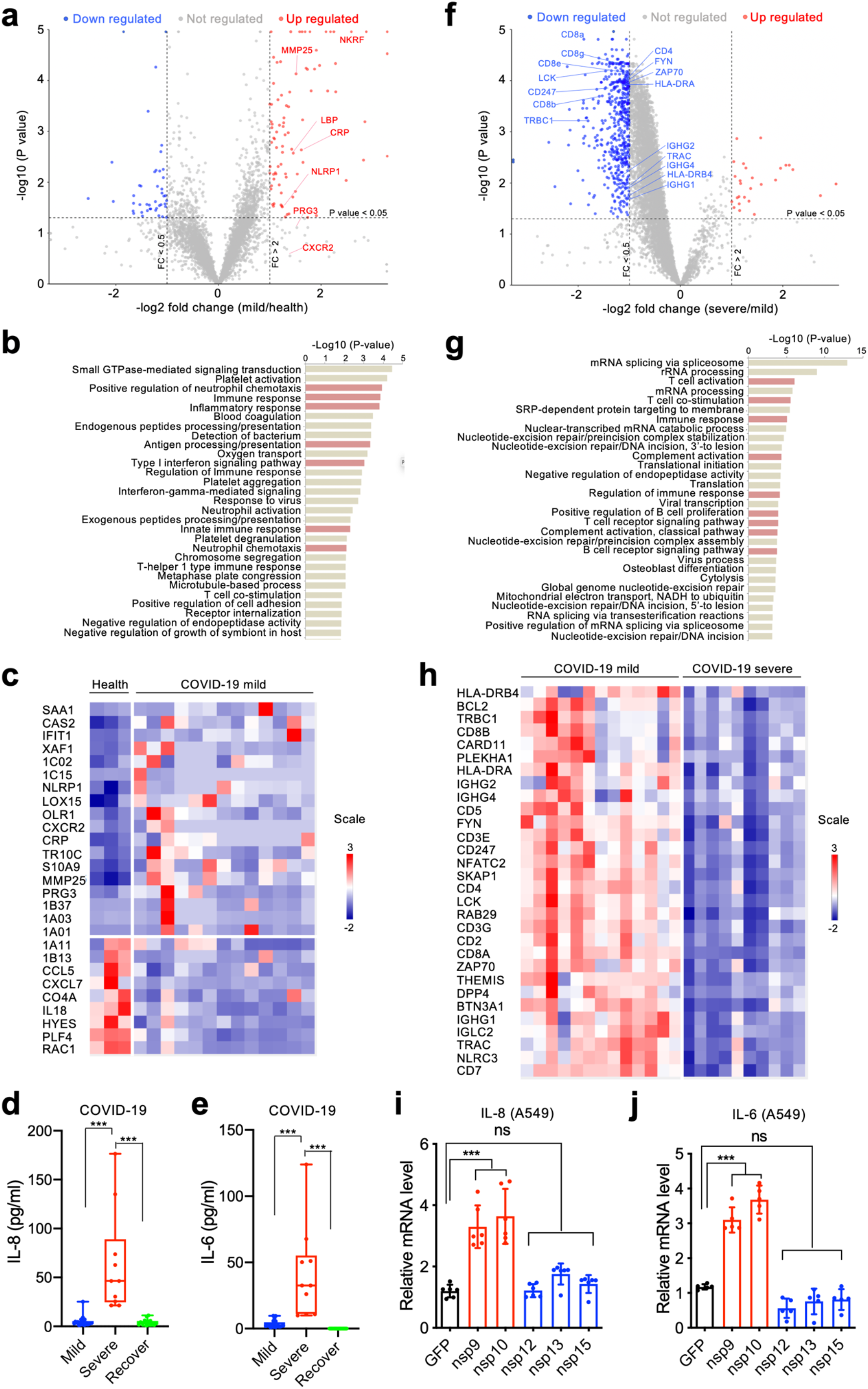
Proteome profile change in PBMCs of COVID-19 patients suggests a potential role of the nsp9/nsp10-NKRF-IL-6/8 axis in neutrophil activation. (**a-c**) Comparative proteomic analysis of COVID-19 patients with mild symptoms (N = 13) compared to healthy control (N = 3). In the volcano plot, each circle represents a protein that was quantifiable in at least three biological replicates. Significantly up- or down-regulated proteins (Adjusted two-sided P≤0.05; |log2(fold change)| ≥ 1) are shown in red and blue, respectively. Differentially expressed proteins related to inflammatory pathways are upregulated and labeled (**a**). Gene Ontology analysis shows the enrichment of biological processes with significantly up- or down-regulated proteins (**b**). Heat map of the differential expressed proteins (|log2(fold change)| ≥ 1) (**c**). (**d-e**) The levels of IL-8 (**d**) and IL-6 (**e**) in PBMCs from COVID-19 patients with mild or severe symptoms or recovered patients (viral RNA negative) were determined by FACS analysis. (**f-h**) Comparative proteomic analysis of COVID-19 patients with severe symptoms (N = 10) compared to mild symptoms (N = 13). Each circle represents a protein that was quantifiable in at least three biological replicates in volcano plot. Significantly up- or down-regulated proteins (Adjusted two-sided P≤0.05; |log2(fold change)| ≥ 1) are shown in red and blue, respectively. Adaptive immunity pathway related differentially expressed proteins are downregulated and labeled (**f**). Gene Ontology analysis shows the enrichment of biological processes with significantly up- or down-regulated proteins (**g**). Heat map of the differential expressed proteins (|log2(fold change)| ≥ 1) (**h**). (**i-j**) Individual stable expression of nsp9 or nsp10 from SARS-CoV-2 significantly promotes the inductions of IL-8 (**i**) and IL-6 (**j**) in the lung epithelial cell line. Mean ± SD; ns, and \*\*\**p* < 0.001 by Student’s *t*-test.

Next, we compared the proteome change between PBMCs isolated from COVID-19-SS and COVID-19-MS patients. Amongst 6,235 quantified proteins (n≥3), 553 proteins were found to be significantly differentially expressed [|log2 (fold change)| ⩾ 1, adjusted two-sided *p* Value ⩽0.05] between COVID-19SS and COVID-19 MS’s PMBCs. 27 of these proteins were upregulated and 526 of these proteins were downregulated in COVID-19-SS as compared to COVID-19-MS (**Fig.3f**, **Extended Data Fig.6a-b** and **Table 3**). Functional analysis by DAVID and Cluster Profiler revealed that these genes are highly enriched in pathways associated with regulation of immune response, T cell co-stimulation and activation, complement activation, B cell receptor signaling pathway, and T cell receptor signaling (**Fig.3g** and **Extended Data Fig.6c-d**). The expressions of T cell receptor (TCR) subunits (CD3e, CD3g, CD247, TRAC, and TRBC1), T cell surface molecules (CD4, CD8a, CD8b, and CD2), MHC class II molecules (HLA-DRA, HLA-DRB1, HLA-DRB4, and HLA-DRB5), T cell migration stimulators (DDP4), and TCR signaling kinases (ZAP70, LCK, and FYN) were dramatically diminished in COVID-19-SS as compared to COVID-19-MS (**Fig.3a,h**), suggesting a failure of T cells activation and function. The expressions of immunoglobulin subunits (IGHG1, IGHG2, IGHG4, IGLC2, IGLL1, and IGHE) were also significant lower in COVID-19-SS (**Fig.3a,h**), indicating reduced antibody secretion by B cells. These results demonstrated that more severe COVID-19 patients undergo a functional decline in adaptive immunity, which is correlated with their lower T cell and B cell populations^11,12^.

### SARS-CoV-2 nsp9 and nsp10 target NKRF to facilitate IL-8/IL-6 production

From the quantitative proteome analysis of PBMCs, it is clear that SARS-CoV-2 induces IL-8/IL-6 expression and increases neutrophil count which may mediate the manifestation of acute respiratory distress syndrome in COVID-19 patients. Therefore, we hypothesized that SARS-CoV-2 virulence factors may potentiate or regulate the induction of IL-8/IL-6 during infection, leading to the uncontrolled infiltration and activation of neutrophils. NKRF is a transcriptional repressor that inhibit IL-8 and IL-6 induction by competing with NF-КB for promoter binding^20,21,29^. From our virus-host interactome, we discovered that NKRF interacts with SARS-CoV-2 nsp9, nsp10, nsp12, nsp13, and nsp15 (**Fig.2** and **Extended Data Fig.7a**). Among these SARS-CoV-2 viral proteins, individually expressed nsp9 or nsp10 facilitated the induction of both IL-8 and IL-6 in lung epithelial A549 cells, while individually expressed nsp12, nsp13, or nsp15 had little or no effects (**Fig. 3i-j**). The levels of TNFα was not affected by either nsp9 or nsp10 (**Extended Data Fig.7b**), suggesting nsp9 and nsp10 does not affect NK-КB signaling transduction overall. Importantly, NKRF deficiency by specific siRNA facilitated the induction of IL-8 and diminished the effect of nsp10, indicating that nsp10 potentially targeted NKRF to regulate IL-8 induction (**Extended Data Fig.7c-d**).

Similar to other coronaviruses, SARS-CoV-2 has underwent rapid adaptive changes against evolutionary pressures since its isolation^30^. Phylogenetic and genetic analysis^31^ of 160 public SARS-CoV-2 sequences reveals 137 amino acid substitutions in 23 viral proteins with no mutations occurring in nsp9 (**Extended Data Fig.7e** and **Table 4**), which indicates nsp9 has remained relatively stable. Unlike nsp9, nsp10 picked up only two amino acid substitutions (A61G and F68L), but with very low variation rate (1/160 for each site) during evolution (**Extended Data Fig.7f**), indicating that nsp9 and nsp10 are potential stable targets to curb SARS-CoV-2-mediated IL-8/IL-6 activation and subsequently recruitment of neutrophils.

Our study integrated virus-host interaction network and quantitative proteome approaches to identify host targets of SARS-CoV-2 viral proteins, demonstrating that SARS-CoV-2 evolved multiple mechanisms to modulate host fundamental cellular processes, which ultimately contributes to its immune evasion and pathogenesis. To our knowledge, this is the first study to characterize the PBMC proteome profile in both mild and severe cases of COVID-19, corroborating the overall clinical features of COVID-19 patients at molecular levels. These systematic analyses not only generate groundwork for future SARS-CoV-2 research on the nature of this virus but also provide valuable targets for specific drug development or repurposing. With these platforms, we identified nsp9 and nsp10 as the potential virulence factors for SARS-CoV-2, which interacts with NKRF to facilitate IL-8/IL-6 induction, ultimately leading to uncontrolled infiltration and activation of neutrophils (**Fig.4**). Specific agents targeting IL-8/IL-6 or the downstream JAK kinases could be repurposed as suitable drug candidates for clinical evaluation. These small molecule inhibitors (i.e. IL-6 or IL-8 receptor antagonists) or agents blocking virulence factor PPIs would have enormous value as anti-virulence therapies in the treatment of SARS-CoV-2 infections (**Fig.4**).

**Figure 4.**
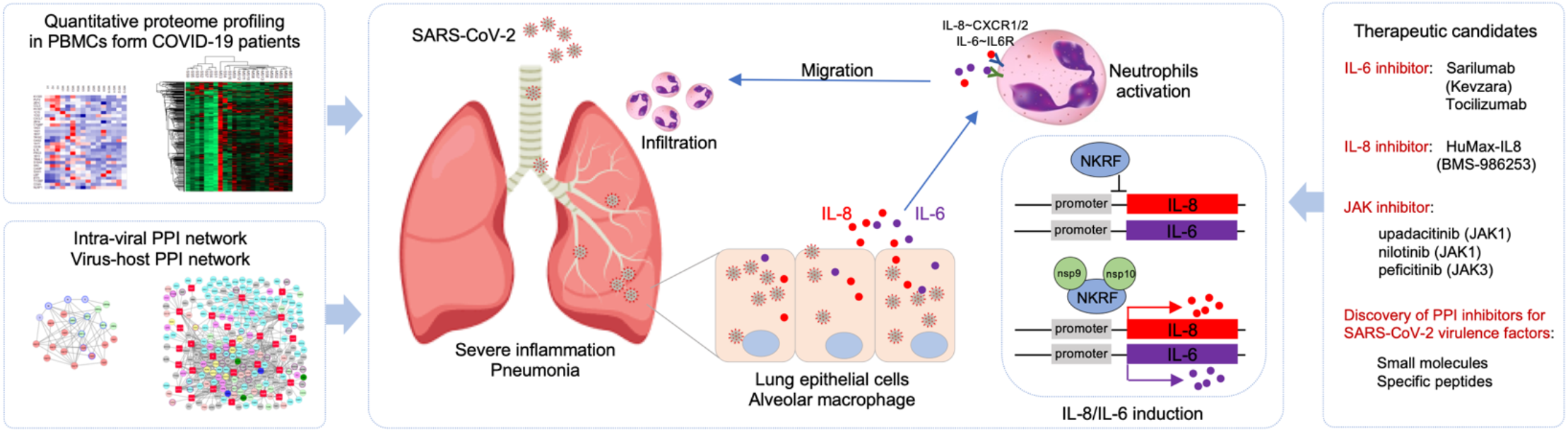
Models of nsp9/nsp10-mediated SARS-CoV-2 pathogenesis and potential design or repurposing for specific inhibitors.

## AUTHOR CONTRIBUTION

Q.L., T.Z., and C.P. conceived of the research, designed the study, and wrote the manuscript. M.G. isolated and lysed PBMCs in denaturing buffer. Z.L., J.L., C.L., X.W., and X.Y. performed the experiments and analyzed data. X.T., P.W., and Y.Y performed the LC-MS analysis. Z.X. helped with visual representation of data and edited the manuscript. All authors commented on the manuscript.

## ACKNOWLEDGMENTS

This work was supported by grants from National Key Research and Development Project of China (2018YFA0900802), National Natural Science Foundation of China (31770176 and 31500667), National Science and Technology Major Project (2020ZX09201001), the Program for Professor of Special Appointment (Eastern Scholar) at Shanghai Institutions of Higher learning, Shanghai Science and Technology Commission (2017QA1403200), Shanghai Municipal Health Commission (2018YQ40 and 201940179), Shanghai Municipal Key Clinical Specialty (shslczdzk01102), Innovative Research Team of High-level Local Universities in Shanghai, the Interdisciplinary Program of Shanghai Jiao Tong University (YG2020YQ14), Chinese Academy of Sciences Large Research Infrastructure of Maintenance and Remolding Project (DSS-WXGZ-2020-0001), and Chinese Academy of Science Key Technology Talent Program.

## METHODS

### Ethics statement

This study was approved by the Ethics Committee of Shanghai Public Health Clinical Center (#YJ-2020-S052-02). The proteomic analysis of PBMC samples were performed on existing samples collected during standard diagnostic tests, posing no extra burden to patients.

### Cell lines, plasmids, and antibodies

HEK293 and A549 cells were maintained in Dulbecco’s modified Eagle’s medium (DMEM; Gibco-BRL) containing 4 mM glutamine and 10% FBS. Vero cells were cultured in DMEM with 5% FBS and 4 mM glutamine. Transient transfections were performed with Lipofectamine 3000 (Thermo Fisher Scientific, #3000015). SARS-CoV-2 genes were synthesized as DNA fragments (Sangon Biotech) and these expression constructs were amplified by PCR and cloned into pLVX-3xFlag-MCS-P2A-tagRFP (puro), pEF-MCS-3xHA, pGBTK7, or pACT2 vectors. Mutations of nsp10 were generated using QuikChange site-directed mutagenesis kit (Stratagene). All constructs were sequenced using an ABI PRISM 377 automatic DNA sequencer to verify 100% correspondence with the original sequence. Primary antibodies were purchased from the following venters: mouse anti-Flag

### Purification of ZIKV viral protein complexes

HEK293 were transfected with 3xFlag-tagged plasmids encoding each SARS-CoV-2 protein by Lipofectamine 3000 (Thermo Fisher Scientific, #L3000015). 3×10^7^ HEK293 cells were harvested in 10 ml lysis buffer (50 mM Tris-HCl [pH 7.5], 150 mM NaCl, 0.5% Nonidet P40, 10% glycerol, phosphatase inhibitors and protease inhibitors) at 72 h post transfection, followed by centrifugation and filtration (0.45 μm) to remove debris. Cell lysates were precleared with 100 μl protein A/G resin, and then incubated with 25 μl anti-Flag M2 resin (Sigma, #F2426) for 12 h at 4°C on a rotator. The resin was washed 5 times and transferred to a spin column with 40 μl of 3 X Flag peptide (Sigma, #F4799) for 1 h at 4°C on a rotator. The purified complexes were sent for mass spectrometry analysis for protein-protein interaction network at the National Facility for Protein Science in Shanghai.

### Virus-host Interaction network analysis

The samples (purified complexes) were precipitated by TCA and resolved in 8 M urea, and then treated with 5 mM TCEP and 10 mM IAA to reduce the disulfide bonds and alkylate the resulting thiol groups, sequentially. The mixture was digested for 16 h at 37 °C by trypsin at an enzyme-to-substrate ratio of 1:50 (wt/wt). The trypsin-digested peptides were desalted by monospin C18 and loaded on a home-made 40 cm-long pulled-tip analytical column (75 μm ID x 360μm OD, ReproSil-Pur C18-AQ 1.9 μm resin, Dr. Maisch GmbH), connected to a Thermo Easy-nLC1000 HPLC system. The samples were analyzed with a 120 min-HPLC gradient from 0 to 100% of buffer B (buffer A: 0.1% formic acid in water; buffer B: 0.1% formic acid in 80% acetonitrile) at 300 nL/min : 4-8 min, 0%-4% B;1-96 min, 8-35% B; 97-104min, 35-60% B; 104-105min, 60-100% B; 105-120min, 100% B. The eluted peptides were ionized and directly introduced into a Q-Exactive mass spectrometer using a nano-spray source with a distal 2.0-kV spray voltage. Survey full-scan MS spectra (from m/z 300-1,800) was acquired in the Orbitrap analyzer with resolution r = 70,000 at m/z 200. One acquisition cycle includes one full-scan MS spectrum followed by top 20 MS/MS events, sequentially generated on the first to the twentieth most intense ions selected from the full MS spectrum at a 28% normalized collision energy. The acquired MS/MS data were analyzed against a UniProtKB Human database (database released on Sept. 30, 2018) containing SARS-CoV-2 viral proteins using Maxquant V1.6.10.43. In order to accurately estimate peptide probabilities and false discovery rates (FDR), we used a decoy database containing the reversed sequences of all the proteins appended to the target database. FDR was set at 0.01. Mass tolerance for precursor ions was set at 20 ppm. Trypsin was defined as cleavage enzyme and the maximal number of missed cleavage sites was set at 3. Carbamidomethylation (+57.02146) of cysteine was considered as a static modification. Methionine oxidation was set as variable modifications.

### Proteomic study of PBMCs from COVID-19 patients

#### Preparation of PMBC cells from patients

The information of 35 patients (13 COVID-19 patients with severe symptoms and 22 COVID-19 patients with mild symptoms) with confirmed SARS-CoV-2 infection and 6 health donors as control was listed in Extended Data Fig.4, including age (47-73), sex, and symptoms. This study followed the Guideline of Novel Coronavirus Laboratory Safety (Edition 2) and was approved by Shanghai Public Health Clinical Center (#YJ-2020-S052-02). The peripheral venous whole blood samples were collected from all the patients and PBMCs were prepared using density-gradient centrifugation followed by washing with cold PBS twice. The cell pellets were lysed by 8 Urea (in 20mM Tris-HCl with protein inhibitor cocktail and phosphor-STOP cocktail, pH 8.0) to denature the proteins and sonicated twice on ice to break the protein-protein or DNA-protein interactions. All the procedures were carried out within the certified laboratory for studies of infectious materials. Then protein samples were combined to generate 3 control samples from 6 healthy donors, 13 mild-case samples from 22 COVID-19-MS and 10 severe-case samples from 13 COVID-19-SS according to the protein concentrations of all the samples. The resulted protein soups were then applied for protein digestion for the following proteomic experiments.

#### Protein Digestion

TCEP (tris(2-carboxyethyl) phosphine, final concentration is 5 mM) (Thermo Scientific) and iodoacetamide (final concentration is 10mM) (Sigma) for reduction and alkylation were added to the solution and incubated at room temperature for 30 minutes, respectively. The protein mixture was diluted four times and digested with LysC at 1:50(w/w) for four hours and then with Trypsin at 1:40 (w/w) (Promega, http://www.promega.com/) overnight.

#### TMT labeling

The concentrations of the peptide soups were measured with Quantitative Colorimetric Peptide Assay (Thermo Scientific). TMT10plex amino reactive reagents (0.8 mg per vial, Thermo Scientific) were suspended in 41 μl of anhydrous acetonitrile and added to peptide soups (100 μg peptides in total). Reactions were allowed to proceed at room temperature for 1 hr, and then quenched by the addition of 8 μl of 5% hydroxylamine for 15 min. The TMT-labeled samples were pooled at a 1:1:1:1:1:1:1:1:1:1 ratio. The mixture was vacuum centrifuged to near dryness and desalted on a MonoSpin™ C18 (GL Science).

#### Off-line high pH-reversed phase fractionation

The desalted TMT-labeled peptides were resuspended in 300 ul 10% buffer C (50 mM ammonium hydroxide, pH 10). Off-line fractionations were performed using a Agilent 1260 HPLC with a 4.6 mm × 250 mm BEH C18, 3.5 μm column (Waters) by a 58 min gradient from 5% to 80% buffer B (100% acetonitrile) with buffer A (100% water) at a flow rate of 0.5 mL/min while buffer C was running constantly at 10% of total flow rate during the process. Fractions were collected over 75 min at 1 min intervals after the start of the gradient in 1.5 ml Eppendorf tubes. The peptide mixture was fractionated into a total of 50 fractions which were consolidated into 18 vials. Samples were subsequently desalted and vacuum centrifuged to near dryness. The desalted peptides were reconstituted in 0.1% fomic acid for downstream LC-MS/MS analysis.

#### LC-MS/MS analysis of peptides

For proteome study, all the 18 fractions for each TMT experiment were analyzed by a home-made 40 cm-long pulled-tip analytical column (75 μm ID x 360μm OD, ReproSil-Pur C18-AQ 1.9 μm resin, Dr. Maisch GmbH), the column was then placed in-line with an Easy-nLC 1200 nano HPLC (Thermo Scientific, San Jose, CA) for mass spectrometry analysis. The analytical column temperature was set at 55 ℃ during the experiments. The mobile phase and elution gradient used for peptide separation were as follows: 0.1% formic acid in water as buffer A and 0.1% formic acid in 80% acetonitrile as buffer B, 0-1 min, 5%-10% B; 1-96 min, 10-30% B; 96-104 min, 30%-50% B, 104-105 min, 50%-100% B, 105-120 min, 100% B. The flow rate was set at 300 nL/min. Data-dependent tandem mass spectrometry (MS/MS) analysis was performed with a Q Exactive Orbitrap mass spectrometer (Thermo Scientific, San Jose, CA). Peptides eluted from the LC column were directly electrosprayed into the mass spectrometer with the application of a distal 2.2-kV spray voltage. A cycle of one full-scan MS spectrum (m/z 300-1800) was acquired followed by top 20 MS/MS events (first mass fixed at 100 m/z scan range), sequentially generated on the first to the twentieth most intense ions selected from the full MS spectrum at a 35% normalized collision energy. Full scan resolution was set to 70,000 with automated gain control (AGC) target of 3e6. MS/MS scan resolution was set to 35,000 with isolation window of 1.8 m/z and AGC target of 1e5. The number of microscans was one for both MS and MS/MS scans and the maximum ion injection time was 50 and 100 ms, respectively. The dynamic exclusion settings used were as follows: charge exclusion, 1 and >8; exclude isotopes, on; and exclusion duration, 30 seconds.

#### Data Analysis

The acquired MS/MS data were analyzed against a UniProtKB human (database released on Sept. 30, 2018) by Maxquant V1.6.10.43. Tolerance of precursor mass and fragment mass were set to ±20ppm. Carbamidomethylation of cysteine (+57.021 Da), TMT-labeled N-terminus and lysine (+229.163 Da) were set as static modifications. Oxidation of methionine (+15.995 Da) was set as variable modification. All identified proteins had a FDR ≤1%, which was calculated at the peptide level.

#### Merging of datasets and statistical analysis

The interpretation of three sets of proteomic results (protein Groups.txt) from MaxQuant were done in the Perseus software environment (version 1.6.2.3) and Python3. Briefly, the datasets were merged by matching the protein accession numbers of each individual dataset for combination by Python in-house developed scripts (comparepg.py). And the protein abundances were normalized by the TMT-labeled internal standard samples which were consisted of an aliquot of each sample and spiked into the 3 sets of TMT labeling experiments. All the normalized reporter ion abundances of biological repeats were averaged for fold-change calculation. For AP-MS analysis, a two-sided two sample t-test was used to obtain the statistical significance to compare the results of the two groups. For proteome analysis, p value was calculated by empirical Bayes moderated t-tests and then adjusted by Benjamini and Hochberg method, using *limma* package^32^ built in R environment. Proteins were required to have a |log2(fold change)|⩾1 and adj. p value cut-off of 0.05 to be considered as being significantly differentially expressed. Gene ontology (GO) analysis was performed using DAVID Bioinformatics Resources 6.8 (https://david.ncifcrf.gov/) and ClusterProfiler v3.14.3^33^.

### Detection of SARS-CoV-2 intra-viral PPIs by yeast-two-hybrid

28 SAR-CoV-2 genes were cloned into both pGBTK7 (bait) and pACT2 (prey) vectors. The reporter Y2HGold yeast strain (Clontech) was co-transformed with different bait-prey combinations as indicated in Extended Data Fig.1 in accordance with instructions for Matchmaker GAL4 Two-Hybrid System 3 (Clontech). The transformed yeast was then screened in 4-dropout plates for the protein-protein interaction.

### RNA extraction and quantitative RT-PCR

Total RNA was isolated from cells with the RNeasy Mini Kit (Qiagen, #74106) and treated with RNase-free DNase according to the manufacturer’s protocol. All gene transcripts were quantified by quantitative PCR using qScript^TM^ One-Step qRT-PCR Kit (Quanta Biosciences, #95057-050) on CFX96 real-time PCR system (Bio-Rad). Primer sequences for qPCR were as follow: IL-6 forward: AACCTGAACCTTCCAAAGATGG, IL-6 reverse: TCTGGCTTGTTCCTCACTACT; IL-8 forward: TCTTGCACAAATATTTGATGC, IL-8 reverse: CCACTGTGCCTTGGTTTC; GAPGH forward: GAGTCAACGGATTTGGTCGT, GAPGH reverse: TTGATTTTGGAGGGATCTCG; TNFα forward: CTCCAGGCGGTGCTTGTTC, TNFα reverse: GGCTACAGGCTTGTCACTCG; NKRF forward: GTAAACATGCAGCTGCCGAC, NKRF reverse: CGTGCACACGGGATTTGAAG.

### siRNA gene silencing

Small interfering RNA (siRNA) targeting human NKRF (#A10001) and negative control siRNAs were purchased from GenePharma. siRNAs were delivered to the A549 cells using Lipofectamine RNAiMAX Transfection Reagent (Thermo Fisher Scientific, #13778030) according to the manufacturer’s instructions. After 48 h of transfection, cells were verified for target gene knock-down by quantitative RT-PCR.

### Phylogenetic and genetic analysis of SARS-CoV-2 sequences

160 SARS-CoV-2 isolates with complete genome sequences from 18 different countries during December 23, 2019 to February 29, 2020 were used for molecular evolutionary analysis in this study. Multiple alignment was performed by Clustal W and phylogenetic tree was constructed by Neighbor-Joining statistical method based on the Maximum Composite Likelihood model with 500 replicates of bootstrap testing using MEGA^31^.

### Quantification and Statistical Analysis

All data were expressed as Mean ± SD as indicated. For parametric analysis, the F test was used to determine the equality of variances between the groups compared; statistical significance across two groups was tested by Student’s t-test; one-way analysis of variance (ANOVA) followed by Bonferroni’s *post hoc* test were used to determine statistically significant differences between multiple groups. *P*-values of less than 0.05 were considered significant.

